# Molecular Transport across Lipid Membranes Controls Cell-Free Expression Level and Dynamics

**DOI:** 10.1101/606863

**Authors:** Patrick M. Caveney, Rosemary M. Dabbs, William T. McClintic, C. Patrick Collier, Michael L. Simpson

## Abstract

Essential steps toward synthetic cell-like systems require controlled transport of molecular species across the boundary between encapsulated expression and the external environment. When molecular species (e.g. small ions, amino acids) required for expression (i.e. expression resources) may cross this boundary, this transport process plays an important role in gene expression dynamics and expression variability. Here we show how the location (encapsulated or external) of the expression resources controls the level and the dynamics of cell-free protein expression confined in permeable lipid vesicles. Regardless of the concentration of encapsulated resources, external resources were essential for protein production. Compared to resource poor external environments, plentiful external resources increased expression by ~7-fold, and rescued expression when internal resources were lacking. Intriguingly, the location of resources and the membrane transport properties dictated expression dynamics in a manner well predicted by a simple transport-expression model. These results suggest membrane engineering as a means for spatio-temporal control of gene expression in cell-free synthetic biology applications and demonstrate a flexible experimental platform to understand the interplay between membrane transport and expression in cellular systems.

## Introduction

Confined cell-free gene expression systems (Karim and Jewett, 2016; Moore et al., 2018; Pardee et al., 2016; Shin and Noireaux, 2012; Siegal-Gaskins et al., 2014; Siuti et al., 2011) are making strides (Perez et al., 2016) toward cell-like capabilities (Scott et al., 2016; Trifonov, 2011). Recent reports demonstrate important steps along this path, including genome replication (Sakatani et al., 2015), metabolism (Garcia et al., 2018), adaptation (Yoshiyama et al., 2018), and growth (Exterkate et al., 2017). However, little work has been reported on one of the key next steps – controlled molecular interactions with the external environment (Collier and Simpson, 2011).

This communication across the membrane is essential for highly complex cellular functions such as chemotaxis (Van Haastert and Devreotes, 2004), symbiosis (Braga et al., 2016), and collective action (e.g. biofilm formation (Flemming et al., 2016)). For cell-like systems, essential next steps like energy harvesting, require the controlled trafficking of molecules across the boundary between cell-like system and the external world. Recent work demonstrates the use of pore-forming proteins (Chalmeau et al., 2011; Noireaux and Libchaber, 2004) or optical treatment of vesicles (Caveney et al., 2019) to engineer transport across lipid membranes. When these transport processes involve molecules that support protein synthesis (e.g. ions, nucleotides, amino acids), transport and gene expression dynamics become entwined (Figure 1A).

**Figure 1.**
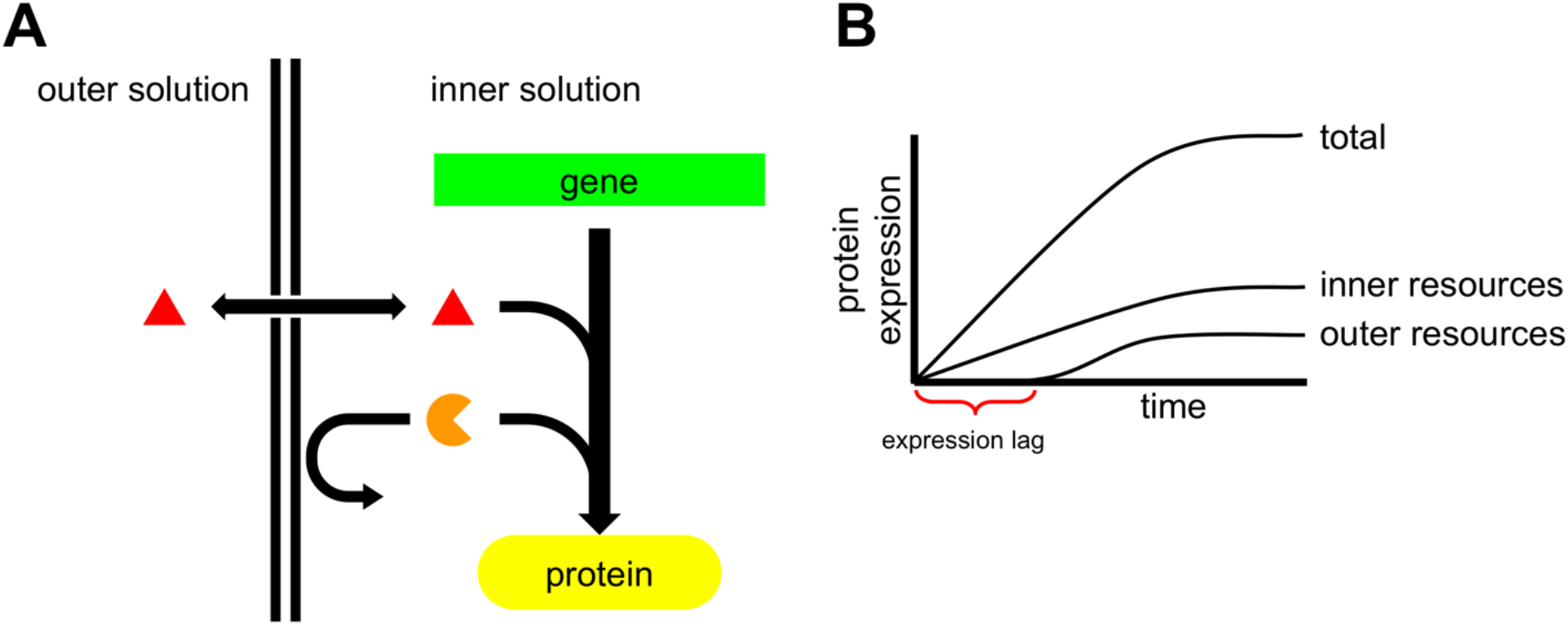
Resource location (internal or external) affects gene expression behavior. (A) Gene expression is affected by the encapsulated (orange circular sector) and external (red triangle) molecular resources. (B) The gene expression transient is the sum of two components: one controlled by the internal resource concentration and one controlled by external resource concentration. The expression transient due to external resources should experience a delay (labeled here as expression lag) related to the membrane transport properties.

Gene expression in cell-like systems is a function of both encapsulated and external expression resources (Figure 1B). Recently, we demonstrated an optical treatment protocol to make vesicle membranes uniformly permeable to molecules relevant to protein expression (Caveney et al., 2019). This technique reduced vesicle-to-vesicle variability in protein expression by making membrane transport properties uniform and by reducing the effects of stochastic seeding of the encapsulated resources. However, permeable membranes change the dynamics of gene expression by accentuating the role of molecular transport across the membrane. Here we show how the location (encapsulated or external) of the expression resources controlled the level and the dynamics of cell-free protein expression confined in permeable lipid vesicles. Regardless of the concentration of encapsulated resources, external resources were essential for protein production, increasing expression levels by ~7-fold. By sourcing essential molecular species, resource-rich external environments rescued expression when internal resources were lacking. Intriguingly, the location of resources and the membrane transport properties dictated expression dynamics in a manner well predicted by a simple transport-expression model. Since the membrane transport properties may be controlled in space and time using a simple optical treatment (Caveney et al., 2019), these results demonstrate the means for predictive spatio-temporal control of gene expression in cell-free synthetic biology applications. Furthermore, the experiments described here show a flexible experimental platform to understand the interplay between membrane transport and expression in individual cells or in groups of cells working cooperatively through cell-to-cell molecular signaling.

## Results and Discussion

To study the dynamics of protein expression in permeable lipid vesicles, we tracked cell-free expression of Yellow Fluorescent Protein (YFP) confined in optically-permeabilized (Caveney et al., 2019) vesicles (Figure 2A and 2B; Methods). Vesicles were created by the emulsion-transfer method described previously (Caveney et al., 2016; Nishimura et al., 2014a; Noireaux and Libchaber, 2004). Briefly, an inner solution composed of the PURE system (Shimizu et al., 2001), a YFP encoding plasmid, pEToppY, and a fluorescent volume marker, AF647 conjugated to transferrin, were vortexed in paraffin oil containing 1-palmitoyl-2-oleoyl-glycero-3-phosphocholine (POPC) to create reverse micelles. The reverse micelles were centrifuged through an oil-water interface into the outer solution to create vesicles (Figure 2A). Vesicles were imaged with z-stacks of 20 slices every 3 minutes for a 3-hour period using a Zeiss, LSM 710 confocal laser scanning microscope. This imaging protocol, light_max, permeabilizes the lipid membranes, allowing the transport of molecular species (e.g. nucleotides, amino acids) essential for expression (Caveney et al., 2019). The images of the AF647 at each time point (Figure 2B, Top) were used to locate individual vesicles possessing distinct boundaries with minimal overlap with neighboring vesicles. The dynamics of protein expression were inferred from the time histories (Figure 2B, Bottom Inset) of the measured YFP fluorescence (Figure 2B, Bottom) from these ROIs.

**Figure 2.**
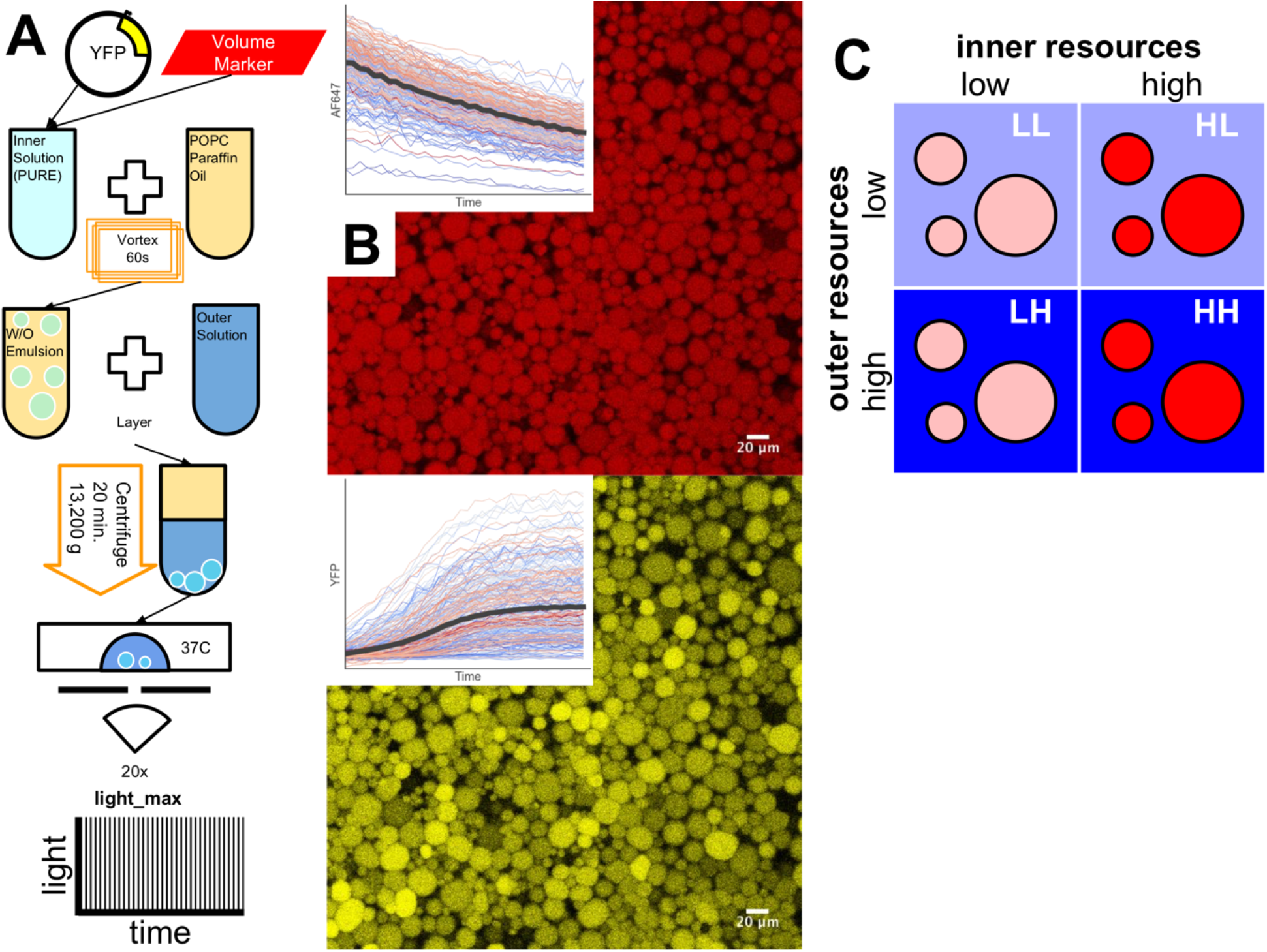
Gene expression in permeable lipid membranes with differing location of expression resources. (A) Method for making and imaging vesicles. Vesicles were imaged using a protocol, light_max, to cause membrane permeability (Caveney et al., 2019). (B) Representative images of the AF647 volume marker and YFP fluorescence in vesicles. Insets show transient behavior of AF647 (photobleaching decay) and YFP fluorescence. (C) Varying the location of essential gene expression resources by changing inner and outer solution concentrations.

We performed experiments using protocols defined by the concentrations (high (H) or low (L); Methods) of resources encapsulated within vesicles and those in the outer solution (Figure 2C). This led to four types of experiments: (1) Low encapsulated/low outer (LL); (2) High encapsulated/low outer (HL); (3) Low encapsulated/high outer (LH); and (4) High encapsulated/high outer (HH). After 2 hours, when protein expression had stopped, the YFP fluorescence resulting from the LL protocol was nearly indistinguishable from background, indicating very low gene expression activity (Figure 3A). Conversely, three of these protocols (HL, LH, and HH) led to significant protein expression activity as indicated by YFP fluorescence levels measurably above background (Figure 3B-D). These results indicate that if the expression resources were available, either encapsulated or within the outer solution, gene expression occurred. Yet, the location of the resources controlled both the level (Figure 3E) and the dynamics (Figure 3F) of expression.

**Figure 3.**
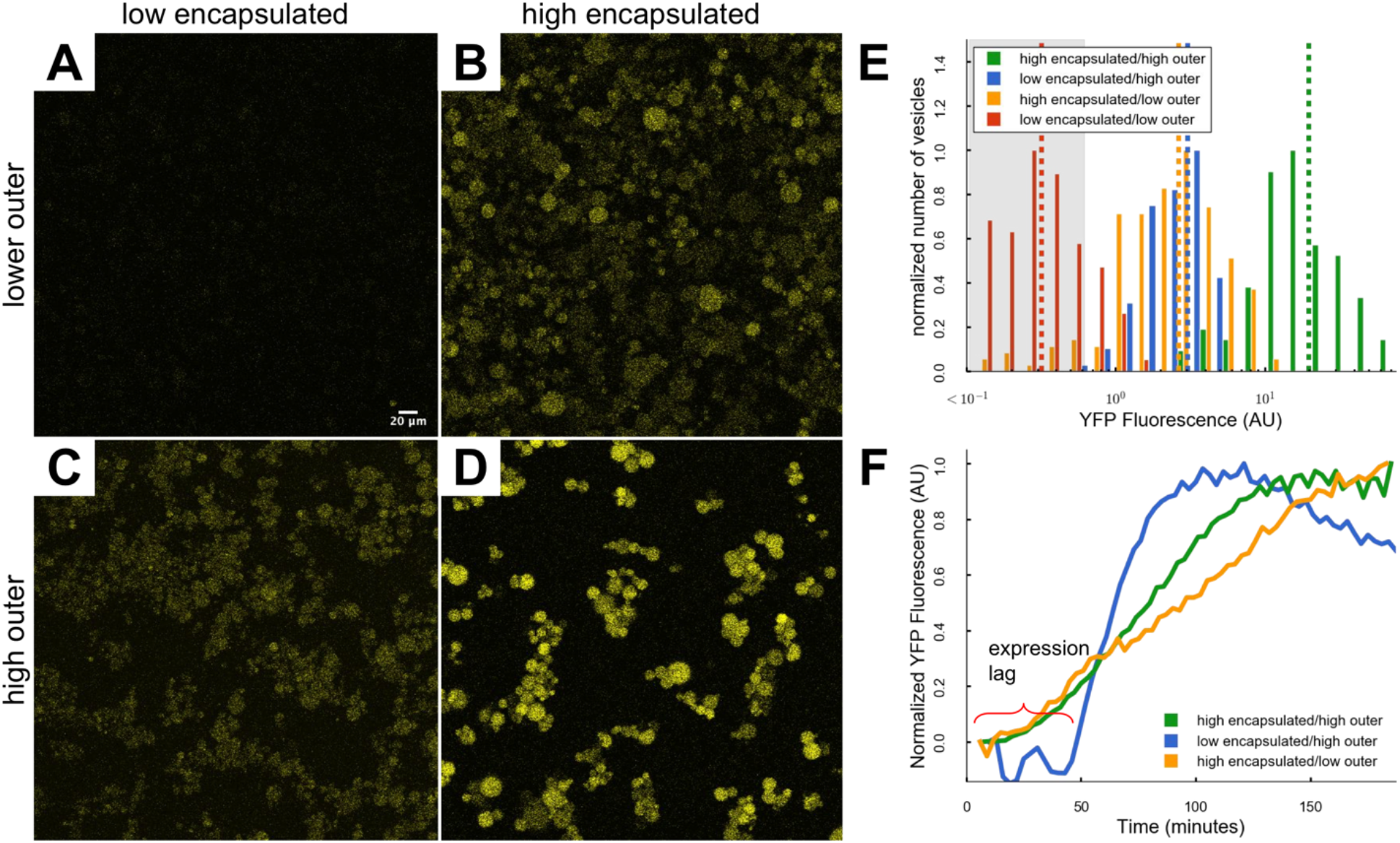
Expression resource location controls gene expression level and dynamics. (A) The t=2 hour image of YFP expressed with the low encapsulated/low outer concentration condition. (B) The t=2 hour image of YFP expressed with the high encapsulated/low outer concentration condition. (C) The t=2 hour image of YFP expressed under the low encapsulated/high outer concentration condition. (D) The t=2 hour image of YFP expressed under the high encapsulated/high outer condition. (E) Distributions of YFP produced under each of the four resource concentration conditions. The dashed vertical lines indicate the means of each distributions. Gray shaded region denotes vesicles with YFP intensities indistinguishable from the background. (F) Average transient behavior of vesicles under the three conditions that produced significant amounts of protein.

With the benefit of a full complement of both encapsulated and external resources, the HH protocol led to the highest level of gene expression (Figure 3D). In agreement with a previous report (Carrara et al., 2018), high resource concentrations were essential to maximizing protein production as expression in the HH condition was nearly ~7-fold greater than either of the other two productive protocols. Intriguingly, HL and LH protocols produced similar levels of total protein at the 2-hour mark (Figure 3B, C, and E), yet displayed very different expression dynamics (Figure 3F). Regardless of the outer solution resource concentration, a high concentration of encapsulated resources led to a rapid onset of expression (Figure 3F). Conversely, expression in the low encapsulated resource environment was rescued by a high concentration of external resources, but only after a considerable delay (Figure 3F).

To better understand the dynamics of cell-free gene expression in permeable vesicles, we constructed a model that accounted for molecular transport across the membrane (Figure 4A). This model included three types of molecular species important in the expression process: (Q) encapsulated molecules (e.g. ribosomes, RNAP) that do not cross the membrane (immobile); (L) highly transportable species (e.g. small ions) that were both encapsulated and in the outer solution that readily crossed the membrane (highly mobile); and (R) slowly transported species (e.g. amino acids, nucleotides) that were both encapsulated and in the outer solution (lowly mobile). In the model, the internal concentration of these three species and the known decay in expression capacity with time (Caveney et al., 2016; Sun et al., 2013) controlled the rate of gene expression. The time dependence of expression capacity was modeled as an exponential decay (rate constant k_p_ = 0.00001*e^-00125*t^). The internal concentration of immobile species was set by the initial conditions (either high or low) and remained constant throughout the simulation (SI Figure 1D). The internal concentrations of the two mobile species (L) and (R) reached equilibrium with the external concentration of these species after a delay time set by the transport rates (SI Figure 1A-C). Since the highly mobile species were expected to equilibrate with the outer solution very quickly, this rate constant was set high (k_Lin/out_=0.1*t). The lowly mobile rate constant was found by fitting to the transport transient of fluorescein into vesicles from the outer solution (Methods, SI Figure 2D). This resulted in time-variant transport process described by k_Rin/out_=0.00019*t, where transport across the membrane for the lowly mobile species increased with increased light exposure.

**Figure 4.**
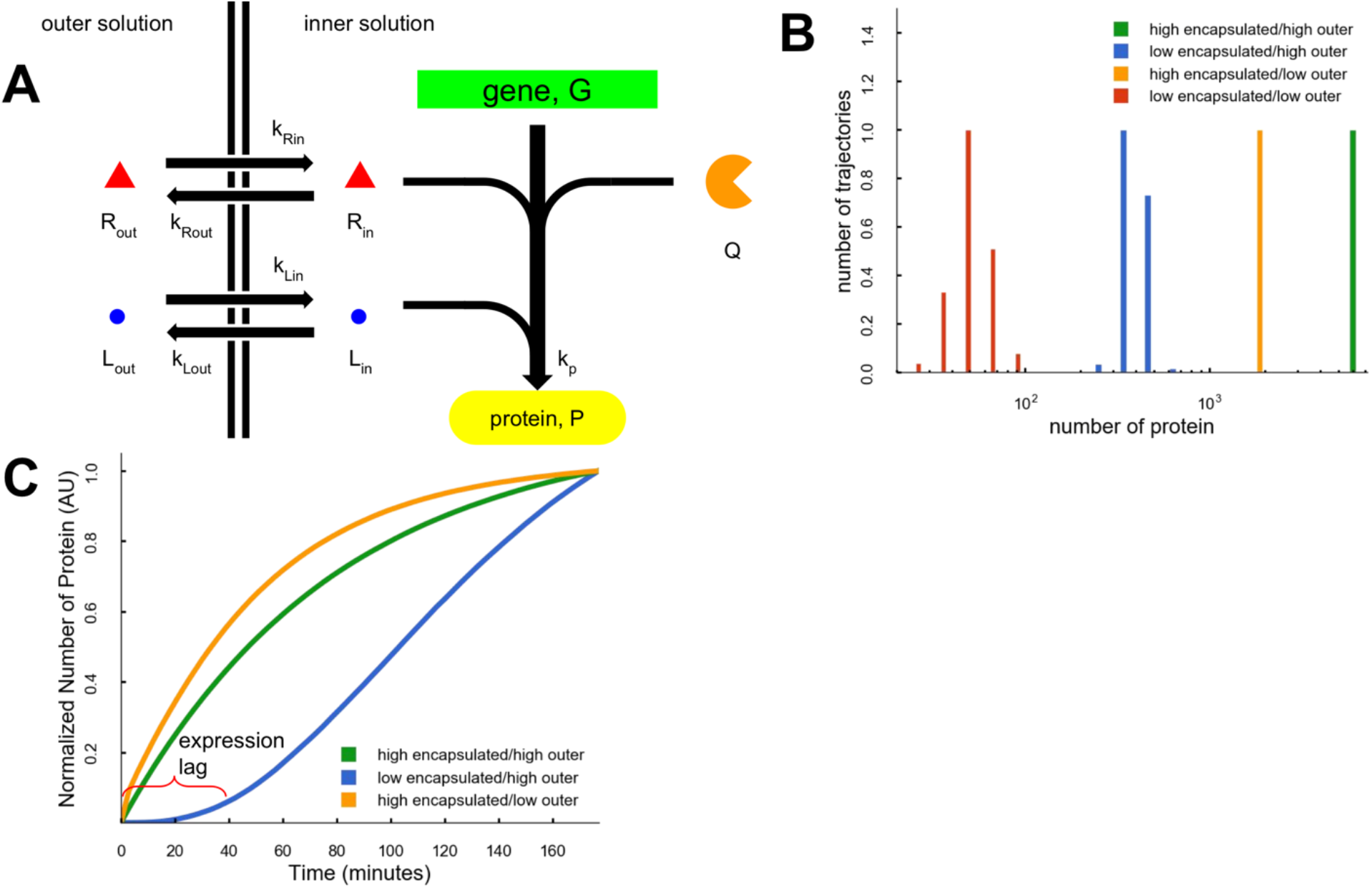
Model of protein expression with resources divided by a permeable membrane. (A) Graphic of the Gillespie simulation. Species G and Q do not cross the membrane while species R and L cross with different rates, k_R_ and k_L_. (B) Distributions of final protein for each of the four conditions. (C) Transients of protein expression for the three conditions that make significant amounts of protein.

Gillespie simulations (Gillespie, 1977) of expression were run using the model for four conditions approximating the experimental HH, HL, LH, and LL protocols, and simulation results reproduced the major experimental behaviors. Similar to experiments, the HH simulations resulted in the most protein. HL and LH simulations resulted in ~3.4x and 14x less protein than HH (Figure 4B). Further, the HH and HL simulations (most encapsulated resources), had a rapid initial rise in protein production while the LH model has a significant time lag (Figure 4C) caused by the delayed arrival of the lowly mobile species. The variation in the amounts of protein made in the different modeling conditions was caused by the interplay between the time that resources arrived in the vesicle and the exponential decay in protein expression capacity.

Perhaps cell and cell-free synthetic biology differ most in the emphasis each place either on gene circuits or on the environment in which gene circuits reside. Cell-free synthetic biology is placing ever greater emphasis on controlling the gene circuit environment by manipulating confinement volume (Caveney et al., 2016), degree of macromolecular crowding (Norred et al., 2018), and the composition of cell extract (Garcia et al., 2018). A defining feature of the gene circuit environment in cells is the controlled transport of molecules across membranes. Such interplay between membrane transport and gene expression in cells play important roles in expression variability (Hansen et al., 2018a) and complex functionality like probabilistic fate determination (Hansen et al., 2018b). Furthermore, cells use membrane transport to take in energy molecules, sense the environment (Van Haastert and Devreotes, 2004), coordinate population behavior (e.g. quorum sensing (Miller and Bassler, 2001)), and share genetic material (e.g. horizontal gene transfer (Gogarten and Townsend, 2005)). It is an intriguing possibility that membrane engineering for controlled molecular transport between encapsulated and external cell-free expression environments may enable essential next steps toward synthetic systems that approach cell-like functional complexity.

## Methods

### Vesicle Preparation

Vesicles were made using the oil-in-water emulsion technique (Noireaux and Libchaber, 2004; Pautot et al., 2003) (Figure 2A). This method encapsulated a protein expressing inner solution in vesicles separated from an osmotically balanced outer solution. The inner solution was prepared using 10 μL Solution A and 7.5 μL Solution B of the PURExpress In Vitro Protein Synthesis Kit from New England Biolabs; 5 μL of sucrose solution (1 M); 0.25 μL of Transferrin-AlexaFluor 647; 0.125 μL of RNAsin (40 U/μL); 0.418 μL (1.67 nM) of YFP encoding pEToppY plasmid (Nishimura et al., 2014b) (200 ng; 478.2 ng/μL stock); and nuclease-free water to bring the total volume of solution to 30 μL. The inner solution was vortexed in 330 μL of paraffin oil containing 30 mg of POPC for 60 seconds. The resulting emulsion was layered above the outer solution and centrifuged at 13,000 g for 20 minutes at room temperature. The low concentration inner solutions were made diluting Solution A and Solution B with nuclease-free water to 1/3 their standard concentrations.

### Outer Solution Preparation

The outer solution for vesicles was mixed from frozen stocks before each experiment. 1.5 μL Amino acid solution, 11.3 μL of ATP (100 mM), 7.5 μL of GTP (100 mM), 0.75 μL of CTP (500 mM), 0.75 μL of UTP (500 mM), 1.8 μL of spermidine (250 mM), 3.75 μL of creatine phosphate (1 M), 4.5 μL of Dithiothreitol (100 mM), 0.75 μL of Folinic Acid (0.5 M), 24 μL of potassium glutamate (3.5 M), 11.3 μL of magnesium acetate (0.5 M), 30 μL of HEPES (1 M), 60 μL of glucose (1 M), and 141.8 μL of autoclaved type I pure water for a total volume of 300 μL. The low concentration outer solutions were made by diluting the entire outer solution to 1/3 its standard concentration.

### Vesicle Imaging

Fluorescent images were obtained using a confocal microscope to track YFP expression and AF647 fluorescence for three hours (Figure 2B and 2B inset). The pellet of vesicles was collected with 100 μL of the outer solution and pipetted onto a no. 1.5 glass bottom petri dish. The lid was placed on the petri dish to minimize airflow and evaporation of the 100 μL outer solution and vesicle drop. Vesicles were imaged with the light_max protocol (Caveney et al., 2019). The petri dish was placed on a Zeiss LSM710 confocal scanning microscope with an incubation chamber warmed to 37°C and imaged every 3 minutes in a z-stack with a 20x air objective. Vesicles were imaged with three lasers: a 405 nm, 6.5 mW laser; YFP was excited with a 488 nm, 6.1 mW laser and fluorescent emission was collected from 515-584 nm; and AF647 was excited with a 633 nm, 1.67 mW laser and fluorescent emission was collected from 638-756 nm. Z-stacks were made of ~21 slices at 1 μm intervals, and the aperture for each slice was 1.00 Airy Units (open enough to allow ~1.5 μm depth of light). The time the vesicles sat on the microscope before imaging was minimized (less than 15 minutes), allowing for imaging for most of the duration of protein expression.

### Data Acquisition and Analysis

Average fluorescent intensities were measured with the FIJI TrackMate (Tinevez et al., 2017) (v3.8.0) plugin. TrackMate found spots with an estimated blob diameter of 10 μm using the Laplacian of Gaussian detector. Spots that were found with an estimated diameter <5 μm, >19 μm, or contrast <0 were removed from the data set. We used the simple Linear Assignment Problem (LAP) tracker to link spots across z-stacks in time to create traces. Traces that had missing frames, traveled >5μm between frames, or tracked for <45 of the 60 frames were removed from the data set. Protein concentration (in arbitrary units) was measured as the average fluorescent intensity for each vesicle at each time point.

### Determining Vesicles Indistinguishable from Background

Background ROIs were determined from the AF647, volume marker, channel. Intensity of these ROIs was measured in the YFP channel. A sample of vesicles with clearly defined edges by eye in the AF647 channel but no clear edges by eye in the YFP channel were measured for their YFP intensity. This value, 0.6147 AU, was slightly above the background value measured and was used as a cut-off for vesicles that were indistinguishable from background fluorescence.

### Measuring the Transient of Membrane Permeability

To investigate the temporal dynamics of membrane transport across the permeabilized membranes, fluorophores were added to the outer solution instead of the inner solution, detailed in (Caveney et al., 2019). Three fluorophores were used, fluorescein (~332 Da), AF633 (~1.2 kDa), and AF647 conjugated to transferrin (~80 kDa), to span the range of protein expression resources, amino acids (~110 Da), nucleotides (~650 Da), proteins (>20 kDa). Fluorescein crossed the membrane in all vesicles by the end of the experiment (SI Figure 2A). The vesicle in the black box is representative. AF633 had vesicles with three behaviors (SI Figure 2B) vesicles in boxes are representative). Behavior (1) vesicles began with high concentrations of AF633 and photobleached. Behavior (2) vesicles started with no AF633 and increased in fluorescence. Behavior (3) vesicles did not increase in fluorescence throughout the experiment. AF647 did not cross the membrane during the experiments (SI Figure 2C). These results show all vesicles are permeable to small molecules but are impermeable to larger molecules.

On average, fluorescence of AF633 and AF647 in vesicles did not increase during the experiment, however fluorescein increased (SI Figure 2D). The fluorescein curve was fit by functions derived from different models of membrane permeabilization. The dashed line assumed first order reaction kinetics, the solid and dotted lines assume permeability increases with time either linearly or sigmoidally. The fluorescein data were better fit by models where permeability increases as the vesicles are exposed to more light.

### Model of Protein Expression in Permeabilized Vesicles

Protein expression inside permeabilized vesicles was modeled with a Gillespie simulation (Gillespie 1977). The model had four populations required to make protein: genes, G, that could not cross the membrane; large resources, Q, that could not cross the membrane; small molecules, R, that could cross the membrane slowly; and very small molecules, L, that could rapidly cross the membrane (SI Figure 1A-D). For all simulations G=10. The initial resource populations, Q, R_in_, L_in_, R_out_, and L_out_, in the high concentration conditions were equal to 100. The initial resource populations, Q, L_in_, R_out_, and L_out_, in the low concentration conditions were equal to 33. The initial R_in_ populations in the low concentration conditions were equal to 0 to model a depletion of inner resources before much protein could be made. The outer solution concentrations, R_out_ and L_out_, were assumed to be constant (SI Figure 3D) because the outer solution volume is much larger than the inner solution volume. Each of the four resource conditions was run for 100 trajectories. Each trajectory was simulated for 180 minutes and sampled every three minutes just as in the experiments.

## Acknowledgements

This research was conducted at the Center for Nanophase Materials Sciences, which is a DOE Office of Science User Facility. P.M.C. and S.E.N. also acknowledge Graduate Fellowships from the Bredesen Center for Interdisciplinary Research and Graduate Education, University of Tennessee, Knoxville. The authors would like to thank Osaka University and Dr. Tetsuya Yomo for providing pEToppYB plasmid, and Dr. Maike Hansen and the Dr. Leor Weinberger laboratory for useful discussions.

## Author Contributions

Conceptualization, P.M.C., C.P.C., and M.L.S.; Methodology, P.M.C., C.P.C., and M.L.S.; Software, P.M.C.; Formal Analysis, P.M.C., M.L.S.; Investigation, P.M.C., R.M.D.; Writing - Original Draft, P.M.C., W.T.M., and M.L.S.; Writing - Reviewing and Editing, P.M.C. and M.L.S.; Visualization, P.M.C.; Supervision, C.P.C., M.L.S.; Funding Acquisition, M.L.S.

