## Supplementary material for "Molecular Transport across Lipid Membranes Controls Cell-Free Expression Level and Dynamics": SI Figure

### Supplemental Figures

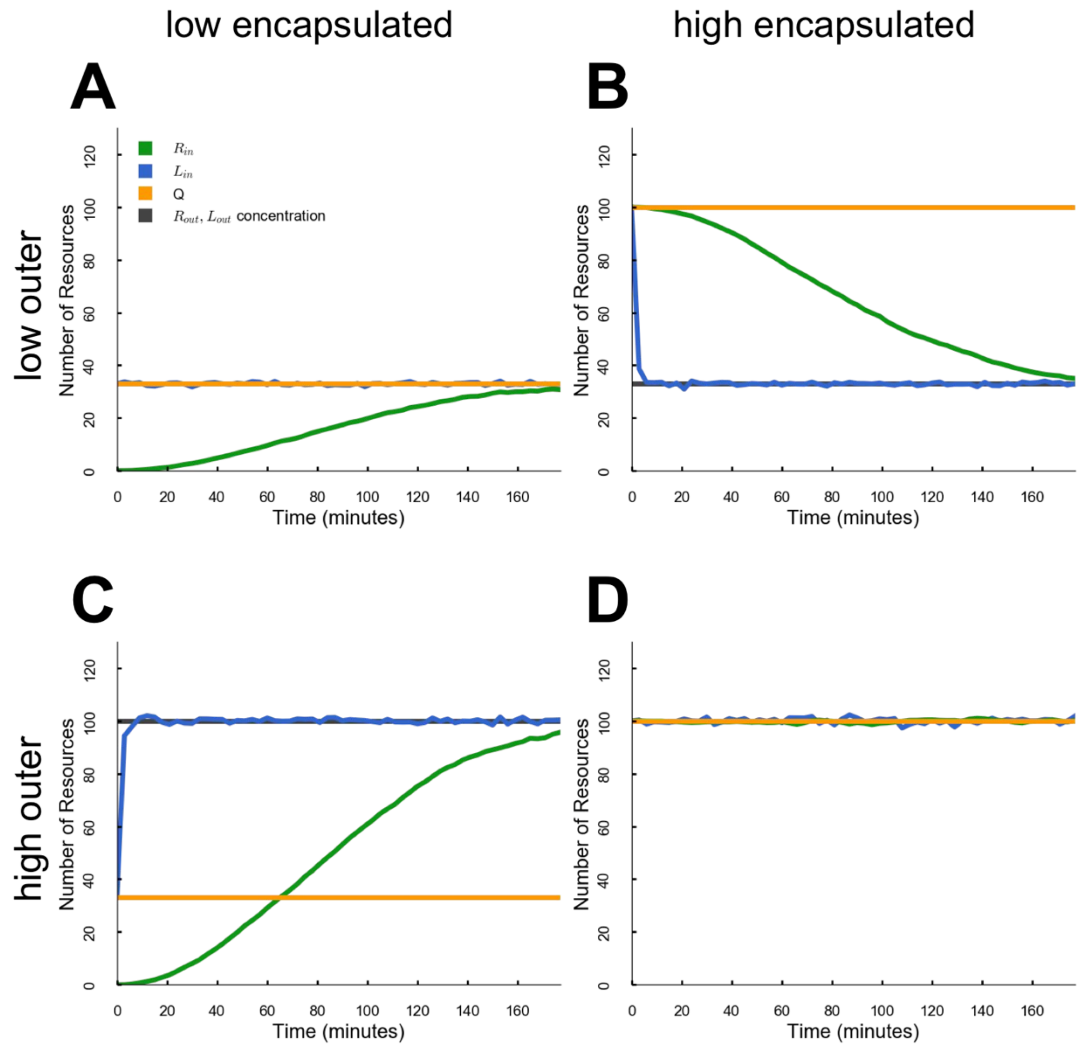


**SI Figure 1: Inner solution resources concentrations in the model.** (A) Behavior of inner resources in the low encapsulated/low outer condition. (B) Behavior of inner resources in the high encapsulated/low outer condition. (C) Behavior of inner resources in the low encapsulated/high outer condition. (D) Behavior of inner resources in the high encapsulated/high outer condition.

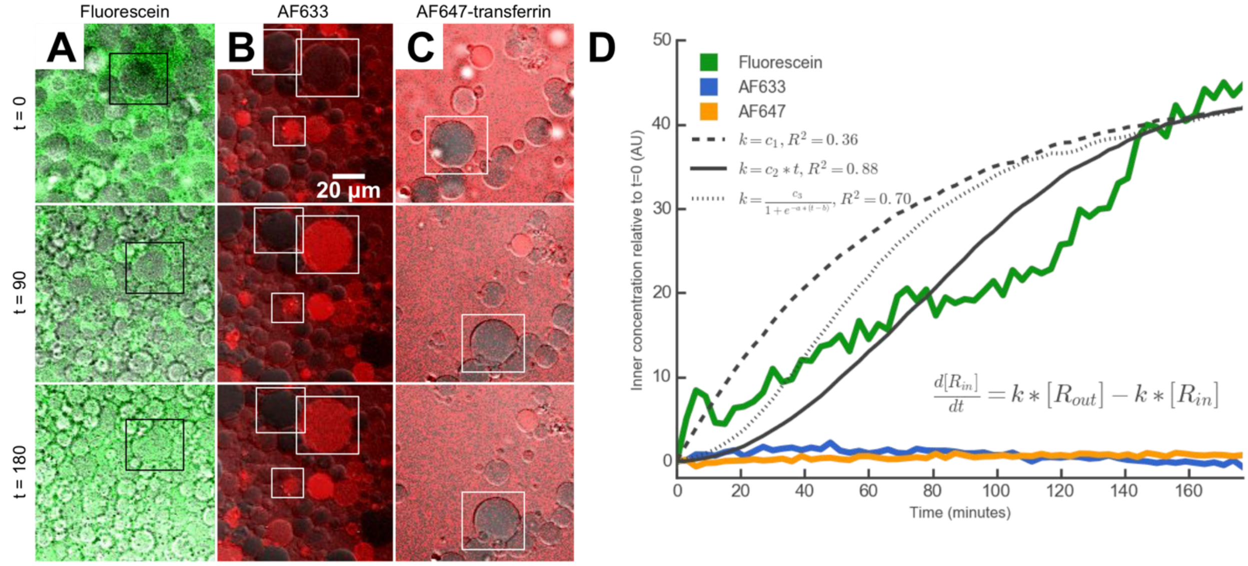


**SI Figure 2: Transient behavior of different fluorophores during to the light_max protocol.** Different fluorophores, (A) Fluorescein, (B) AF633, and (C) AF647, were added to the outer solution and imaged with the light_max protocol. At t = 0 minutes most vesicles in each case have a lower intensity than the outer solution. At t = 180 minutes, (A) fluorescein in all the vesicles has equilibrated with the outer solution while (B) AF633 and (C) AF647 mostly remained lower intensity than the outer solution. (D) Average fluorescent intensity transients for the three fluorophores during the light_max protocol. Gray lines are the $[R_{in}]$ population of the LH model resulting from three different types of membrane permeabilization fit to the fluorescein curve.
